# Deciphering temporal antifungal dynamics of a rare actinomycete via integrated omics

**DOI:** 10.64898/2026.02.10.705106

**Authors:** Hildah Amutuhaire, Michael Dubovis, Ivan Plyushchenko, Judith Kraut-Cohen, Tal Luzzatto-Knaan, Jonathan Friedman, Eddie Cytryn

## Abstract

Fungal phytopathogens pose a persistent threat to global crop production, and widespread use of chemical fungicides has driven resistance development and environmental concerns, necessitating sustainable alternatives. Actinomycetes produce diverse bioactive metabolites, yet natural product discovery has disproportionately focused on *Streptomyces*, leaving rare actinomycete taxa underexplored. *Saccharomonospora xinjiangensis* XJ-54 is a rare actinomycete exhibiting strong antifungal activity against Fusarium phytopathogens, including *Fusarium oxysporum f. sp. cucumerinum* (FORC), and harbors numerous biosynthetic gene clusters (BGCs) of unknown function. However, as in many rare actinomycetes, BGCs may be transcriptionally silent under standard laboratory conditions, and their expression dynamics remain poorly understood. To elucidate the molecular basis of antifungal activity in *S. xinjiangensis* XJ-54, we integrated genomic, transcriptomic, and metabolomic analyses. Cell-free supernatants inhibited FORC after 5 days of fermentation, with activity increasing by day 7. Time-resolved RNA sequencing demonstrated that all genomically-identified BGCs were transcriptionally active but exhibited distinct growth-phase-dependent expression patterns, with approximately half upregulated during exponential growth, and the remainder following transition to stationary phase. We observed temporal variations in transcriptional coupling between cluster-specific regulators and biosynthetic genes. LC-MS-based metabolomics showed growth-phase-dependent metabolite shifts, including stationary phase accumulation of secoiridoid-like monoterpenoids, N-acyl amines, and alkaloids (imidazoles, pyridines, indoles), correlating with the observed antifungal phenotype. Bioactivity-guided fractionation subsequently yielded an active fraction containing a predicted halogenated alkaloid that induced hyphal damage in FORC. These findings indicate that antifungal activity in *S. xinjiangensis* XJ-54 arises from temporally coordinated biosynthetic programs, providing a framework for optimizing growth conditions and prioritizing BGCs for functional characterization.

**Importance:** The discovery of new antifungal compounds is critical for sustainable agriculture and medicine, yet most natural product research has focused on screening a narrow range of well-studied microorganisms. Rare actinomycetes represent an untapped reservoir of chemical diversity, but their biosynthetic potential is predominantly unknown. By integrating time-resolved transcriptomics with metabolomics, we show that the rare actinomycete *Saccharomonospora xinjiangensis* XJ-54 produces antifungal metabolites through temporally coordinated biosynthetic programs. Contrary to the prevailing assumption that the majority of biosynthetic gene clusters (BGCs) are silent, all BGCs in this strain were transcriptionally active under standard cultivation conditions, with expression patterns that were strongly growth-phase dependent. This work provides a roadmap for unlocking the biosynthetic potential of rare actinomycetes and accelerating the discovery of antifungal natural products that can be applied in agriculture and human health.

## Introduction

Plant fungal diseases threaten global food security, causing over 20% annual crop losses despite intensive fungicide use (1, 2). Fusarium species are among the most destructive phytopathogens, causing devastating wilt, rot, and blight diseases across major crops. The decline in fungicide efficacy due to resistance development, coupled with mounting environmental and health concerns, has intensified the search for biological alternatives (3–5). Soil-derived bacteria synthesize a diverse repertoire of secondary metabolites (SMs) that, while not essential for growth, mediate critical ecological interactions including pathogen antagonism (6, 7). Bacterial SMs are therefore a prolific source of antifungal compounds for sustainable crop protection, and for clinical applications, where effective treatment options are increasingly limited by rising resistance and a severe shortage of new therapeutics (8).

Advances in genome sequencing and bioinformatics-driven mining have unveiled an extensive reservoir of diverse SM-encoding biosynthetic gene clusters (BGCs) within microbial genomes particularly actinomycetes, underscoring their vast metabolic potential (9). Despite this potential, natural product discovery has remained disproportionately focused on *Streptomyces*, a single genus within actinomycetes, resulting in repeated rediscovery of known compounds (10, 11). In contrast, rare actinomycete lineages possess genomes of comparable complexity with a diverse array of systematically underexplored BGCs, whose products and ecological roles are largely unknown (11). Actinomycetes are highly abundant in soil and rhizosphere, and it is postulated that strains isolated from disease-suppressive soils possess antifungal biosynthetic machinery, making them an ideal source for discovering antifungal compounds from underexplored taxa (12).

A critical bottleneck in natural product discovery is the difficulty in reliably linking BGCs to their cognate metabolites, largely because many BGCs are transcriptionally silent under standard laboratory conditions and require specific regulatory or environmental cues for activation (13, 14). Furthermore, predicting which BGCs will be expressed, and at levels sufficient to yield detectable metabolites under given cultivation conditions, remains highly uncertain (15). These constraints highlight the need for integrated multi-omics approaches that couple transcriptomics and untargeted metabolomics, which link gene expression and chemical signatures enabling identification of transcriptionally active clusters, and providing insights regarding the dynamics of metabolite production (16–18). Such integrative analyses are particularly valuable in poorly characterized taxa, where reference compound databases and known regulatory models are lacking.

A recent shotgun metagenomic investigation of compost-amended cucumber rhizospheres revealed *Saccharomonospora* among the differentially abundant taxa associated with suppression of *Fusarium oxysporum* f. sp. *cucumerinum* (FORC) (19), suggesting involvement in pathogen antagonism. This ecological evidence prompted us to investigate the functional potential of *Saccharomonospora*, an underexplored actinomycete genus. We screened multiple strains for antagonism against a panel of phytopathogenic Fusarium species, and among these, *Saccharomonospora xinjiangensis* XJ-54 exhibited the most pronounced and broad-spectrum antifungal activity (20). In addition, genome analysis of this strain revealed numerous BGCs with no detectable homology to known clusters, suggesting the capacity to produce previously uncharacterized metabolites.

In this study, we evaluated and optimized culture conditions for *S. xinjiangensis* XJ-54 to maximize the antifungal activity of its extracellular metabolites. We simultaneously integrated transcriptomic and untargeted metabolomic analyses to elucidate the temporal dynamics of BGC expression and metabolite production relative to observed antagonistic phenotypes. Bioassay-guided fractionation identified highly active excreted fractions corresponding to expressed BGCs, uncovering a platform for the discovery and evaluation of antifungal metabolites with potential applications in public health and agriculture.

## Materials and Methods

### Bacterial strain cultivation

*S. xinjiangensis* XJ-54 (DSM 44391) was obtained from the Leibniz Institute DSMZ-German Collection of Microorganisms and Cell Cultures GmbH (Braunschweig, Germany; https://www.dsmz.de) (21). The strain was cultured in tryptic soy broth (TSB; BD, USA) or on tryptic soy agar (TSA) and preserved in 25% (v/v) glycerol at −80 °C. For routine use, frozen stocks were streaked onto TSA plates and incubated at 30 °C for 7 days to allow colony maturation. Liquid starter cultures were prepared by inoculating 5 mL of TSB with cells scraped from TSA plates and incubating at 30 °C with shaking at 170 rpm for 3-4 days.

To determine bacterial growth rate, 1 mL of the starter culture was used to inoculate triplicate flasks containing 50 mL of fresh TSB. Optical density at 600 nm was measured every 24 h for 7 days, with three readings averaged per replicate. The exponential phase was identified between 48 h and 96 h, and the stationary phase between 96 h and 168 h. Based on this growth curve, three time points were selected for antifungal, transcriptomic, and metabolomics analyses: 72 h (exponential phase), 120 h (early stationary phase), and 168 h (late stationary phase).

### Fungal cultivation

The GFP-labeled FORC strain (22) was used for in vitro antifungal activity assays as described below. Cultures were stored in 50% glycerol at -80°C and maintained on potato dextrose agar (PDA; BD Difco Laboratories, Detroit, MI, USA) supplemented with 250 mg/L chloramphenicol (hereafter referred to as PDA+; Thermo Fisher Scientific, Waltham, MA, USA) at 25°C. GFP-FORC was initially thawed on PDA supplemented with 100 μg/mL hygromycin (Thermo Fisher Scientific) and then cultured on PDA+ for routine cultivation. Spore solution was prepared by inoculating a mycelial plug from a 5-day-old PDA+ plate into 25 mL of fresh potato dextrose broth (PDB) in a 250-mL Erlenmeyer flask. Flasks were incubated for 7 days at 25°C with shaking at 100 rpm. Spores were extracted by filtering the PDB culture through a 40-μm nylon cell strainer (Corning Inc., Corning, NY, USA), counted using a hemocytometer, and diluted in PDB to 1 × 10⁵ spores/mL.

### Evaluation and optimization of antifungal activity of *S. xinjiangensis* XJ-54

To evaluate the antifungal activity of *S. xinjiangensis* XJ-54 extracellular metabolites, FORC spores were exposed to cell-free supernatants (CFS) as previously described (19). Briefly, 1 mL of 3-day-old XJ-54 starter culture was inoculated into 50 mL TSB (BD Bacto™ Soybean-Casein Digest Medium) in 250 mL flasks and incubated at 30°C with shaking (170 rpm). Triplicate flasks were prepared for independent sampling at 3, 5, and 7 days. After incubation, cultures were centrifuged (5000 × g, 10 min), and supernatants were filtered (0.22 µm Millex-HV; Sigma-Aldrich, Israel) before use in antifungal assays.

To assess the influence of incubation temperature on antifungal activity, *S. xinjiangensis* XJ-54 was cultured at 25°C (below optimal), 30°C (optimal), and 40°C (above optimal) for 7 days. To obtain crude extracts, CFS were subjected to C18 solid-phase extraction (Discovery® DSC-18 SPE Tube, 52604-U; Supelco, Sigma-Aldrich, Israel). For identification of solvents that efficiently extract antifungal metabolites, samples were eluted with either 100% methanol or 100% acetonitrile (both HPLC-grade), which differ in polarity and enable selective extraction of distinct metabolite classes. Eluates were evaporated to dryness (speed vacuum, 25°C), resuspended in 1 mL sterile distilled water, filtered (0.22 μm), and tested for antifungal activity as described below.

All antifungal assays were performed in vitro by transferring 100 μL of FORC spore suspension (1 × 10⁵ spores/mL) into each well of a 96-well plate (Biosigma, Venice, Italy), followed by addition of 100 μL CFS or 50 μL crude extracts. Treated spores were incubated at 25°C for 24 hours, and growth was monitored by measuring fluorescence (488/528 nm) and optical density (600 nm) using a multimode microplate reader (Synergy H1, BioTek, USA). Cycloheximide (Sigma-Aldrich) and TSB served as positive and negative controls, respectively.

### Assessment of fungal cell viability and cell wall integrity

Viability of FORC cells treated with crude extract fractions was evaluated using a previously described propidium iodide (PI) based protocol (23), and calcofluor white (CW) was used to visualize fungal cell walls and assess structural integrity. FORC spore suspensions (200 μL, 1 × 10⁵ spores/mL) were treated with 10 μL of test fractions (resuspended in DMSO, final concentration 0.05%) for 18 h. Samples were centrifuged at 4000 rpm, 10 min, 4°C, resuspended in 200 μL sterile saline, and stained with 0.2 μL PI (20 mM; L7012 LIVE/DEAD® BacLight Kit, Invitrogen, Waltham, MA, USA). The samples were kept in the dark for 30 min at room temperature and immediately before imaging, at which 0.5 μL CW was added. Ten microliters of the stained suspensions were observed using a Leica SP8 confocal laser scanning microscope with 488 and 552 nm lasers and a 40× objective (HC PL APO CS, Leica, Wetzlar, Germany). Images were processed using Leica Application Suite X software. Membrane and cell wall integrity were assessed by examining red PI staining (indicating membrane damage) and CW staining patterns. Three replicates were used per treatment.

### DNA extraction, genome sequencing and annotation

Although *S. xinjiangensis* XJ-54 has been previously sequenced using short-read methods (NCBI bioproject: PRJNA62221), we re-sequenced the genome using long-read technology and performed a hybrid assembly to generate a fully closed genome. High molecular weight genomic DNA was isolated from *S. xinjiangensis* XJ-54 using the MagAttract HMW DNA extraction kit (Qiagen, USA) following the manufacturer’s protocol. DNA quality and quantity were assessed using a NanoDrop spectrophotometer, Qubit fluorometer, and agarose gel electrophoresis to ensure high integrity for long-read sequencing. Whole-genome sequencing was performed by Plasmidsaurus Inc. (Eugene, USA) using Oxford Nanopore Technologies (ONT) and Illumina sequencing platforms. Long-read assemblies were generated using Canu (v2.1.1) (24), Flye (v2.6) (25), and Raven (v1.1.10) (26), and integrated using Trycycler (v0.3.1) (27) to produce a consensus genome assembly. The resulting draft assembly was polished with Medaka (v1.4.4) using ONT reads, followed by short-read error correction using Polypolish (v0.6.0) (28) with Illumina data. Genome completeness and contamination were evaluated using checkM2 (v1.0.2) (29), functional annotation performed with Bakta v1.9.2 (30), and BGCs were predicted using antiSMASH v7.1.0 (31). The final genome assembly has been deposited in GenBank under PRJNA1397885 and complements the previously available reference genome PRJNA62221.

### RNA Extraction, Sequencing, and annotation

Total RNA was extracted from 2 mL of culture at each timepoint using the NucleoSpin® RNA Isolation Kit (Macherey-Nagel, Düren, Germany), following the manufacturer’s instructions. RNA samples were sent to the Genomics and Microbiome Core Facility at Rush University (Chicago, USA) for sequencing. Libraries were prepared using the Zymo-Seq RiboFree Total RNA Library Kit (Zymo Research) for rRNA depletion and cDNA synthesis, and sequencing was performed on the Illumina NovaSeq X platform (2 × 150 bp paired-end) according to the facility’s standard protocols. The quality of raw sequencing reads was assessed using FastQC. Adapter trimming and quality filtering were carried out using BBduk (https://sourceforge.net/projects/bbmap/), employing the Phred trimming method at Q10 and removing reads shorter than 45 bp. High-quality reads were aligned to the *S. xinjiangensis* XJ-54 reference genome using Bowtie2 (32), and alignments were processed with SAMtools (33). Gene-level quantification was performed using FeatureCounts (34). Transcript counts were normalized to transcripts per million (TPM) using the EdgeR package (35) and expression values for BGCs were calculated as the mean TPM of core biosynthetic genes annotated by antiSMASH.

Differential expression analysis across the analyzed timepoints was conducted using DESeq2 (36). K-means clustering (k = 5) was performed in R, using z-score-transformed TPM values to group genes with similar expression dynamics across time points. Protein-coding genes were annotated using KOfamScan for KEGG Orthology (KO) assignment and RPS-BLAST against the NCBI Conserved Domain Database for classification into Clusters of Orthologous Groups (COGs). Enrichment analysis of differentially expressed genes in KEGG pathways was performed using KOBAS 3.0 (37). The raw transcriptomic data have been deposited in the NCBI Sequence Read Archive under BioProject accession number PRJNA1397920.

### Untargeted metabolomics analysis

Chromatographic separation was performed using a Vanquish UHPLC system (Thermo Scientific) equipped with a Kinetex C18 reversed-phase column (1.7 µm, 50 × 2.1 mm, 100 Å pore size; Phenomenex). The mobile phase consisted of water with 0.1% formic acid (A) and acetonitrile with 0.1% formic acid (B), delivered at a constant flow rate of 0.3 mL/min. The gradient program was as follows: held 1 min at 5% B, then to 95% B over 7 minutes, held at 95% B for 2 minutes, decreased to 5% B within 1 minute, and held at 5% B for 4 minutes to re-equilibrate the column. The MS analysis was performed on a *tims*TOFPro2 mass spectrometer (Bruker Daltonics), controlled by *tims*Control and Hystar software packages (Bruker Daltonics), and equipped with ESI source operating in positive or negative ion mode. External mass calibration was performed using sodium formate. MS1 spectra were acquired over an m/z range of 50–2000 at a scan rate of 1 Hz. Data-dependent MS/MS (Auto MS/MS) acquisition was triggered at a rate of up to 12 Hz to collect fragmentation spectra for precursor ions. Raw data files were converted to mzML format and used for downstream processing as in the sections described below.

### Molecular networking and spectral annotation

Feature-based molecular networking (FBMN) was carried out using the GNPS platform (https://gnps.ucsd.edu) (38, 39) Prior to network generation, mass spectrometry data were processed in MZMINE 4.6.1 (40) to perform peak detection, deconvolution, alignment, and feature quantification. The exported results were used as input for FBMN. The data was filtered to remove fragment ions within ±17 Da of the precursor m/z. Additionally, MS/MS spectra were window-filtered by retaining only the six most intense fragments in a ±50 Da window across the spectrum. The precursor and fragment ion mass tolerances were each set to 0.02 Da. A molecular network was built by connecting nodes with cosine scores greater than 0.7 and at least six matched peaks. Edges were retained only when both connected nodes occurred in each other’s top 10 most similar spectra. Molecular families were restricted to a maximum size of 100 nodes by iteratively removing the lowest scoring edges. Spectral library matching was performed against GNPS spectral libraries (38, 41), using the same filtering parameters. Only matches with a cosine score above 0.7 and at least six matching fragment ions were retained. The MS/MS spectra was further annotated using DEREPLICATOR (42). The MS/MS fragmentation spectra was also analyzed using SIRIUS version 6.1.0 (43) for molecular formula identification and CSI:FingerID (44) for structure database search and substructure annotation and CANOPUS (45) for *de novo* compound class prediction. Network visualization was done using Cytoscape (46). The molecular networking job can be publicly accessed at https://gnps.ucsd.edu/ProteoSAFe/status.jsp?task=4f3d9c6461f34bcca1c10f3107ff22a4. Peak lists derived from the aligned peaks and associated spectra served as input for statistical analyses conducted in MetaboAnalyst6.0 (47) and MaAsLin2 (48).

### Fractionation of bioactive extracts

Crude extracts from 7-day cultures were pooled and fractionated by flash chromatography using a Teledyne ISCO Combiflash RF-300 automated system. Approximately 2 mg of dried sample was dissolved in 1 mL of acetonitrile and loaded on Redisep Column (RediSep R_f_ 30g C18 High Performance Gold Column, 69-2203-335, TELEDYNE ISCO). The spectrum was monitored with the following wavelengths: λ_1_ 214 nm; λ_2_ 254 nm and with a total run time of 10.7 min. The mobile phase consisted of water with 0.1% formic acid (A) and acetonitrile with 0.1% formic acid (B). The gradient employed for preparative separation was 10% of B held for 0.4 min, 7.1 to 100% B in 6.7 min, held at 100% B for 1.8 min, 100 to 50% B in 0 min and kept at 50% B for 1.4 min at a flow rate 40 mL/min throughout the run. Fractions were collected in 18 mL tube and tested for antifungal activity as described above.

#### Data visualization

Figures were generated using *ggplot2* (v3.5.2; (49)) in R (v4.0.3; RStudio v2025.09.2 Build 418, Windows) and GraphPad Prism.

## Results

### Genome sequencing and secondary metabolite biosynthetic potential

To provide a high-confidence genomic reference for downstream multi-omics analyses, a high-quality *de novo* assembly of *S. xinjiangensis* XJ-54 was generated from long-read and short-read sequences. The final assembly consisted of a single circular chromosome spanning 4.77 Mb with 69.0% GC content, 90.6% coding density, and 4,342 predicted protein-coding sequences, with 99.99% completeness and 0.70% contamination. Using antiSMASH v7.1.0, we identified 12 BGCs encompassing ∼9% (401 genes) of the coding genome, none of which were contig edge-associated, indicating complete cluster boundaries. Only one BGC (Region 10, encoding ectoine) shared >50% sequence similarity with characterized clusters in the MIBiG database (**Table S1**), while the remaining 11 showed minimal similarity (<50%), highlighting the biosynthetic novelty of this strain.

### Stationary-phase and cultivation at 30 °C enhance *S. xinjiangensis* XJ-54 antifungal metabolite production

To determine the growth conditions that enhance antifungal activity against FORC, we monitored the growth dynamics of strain *S. xinjiangensis* XJ-54 in TSB medium and evaluated its CFS for inhibitory activity across different growth phases and temperatures. The strain displayed a clear growth progression, entering mid-exponential phase after approximately 3 days and reaching stationary phase between 5 and 7 days (Figure 1A). CFS collected from cultures incubated at 30 °C for 3, 5, and 7 days was tested for antifungal activity by monitoring the growth of CFS-treated FORC spores over 24 hours. While 3-day CFS did not inhibit FORC, significant growth inhibition was observed with 5-day CFS, and the antifungal effect increased further with 7-day CFS (**Figure 1B, C**). To assess the effect of temperature, antifungal activity of CFS from 7-day cultures grown at 25 °C, 30°C, and 40 °C was compared. Inhibitory activity was significantly higher in CFS of cultures grown at 30 °C compared to 25 °C and 40 °C (**Figure 1D, E**). These findings revealed that stationary-phase cultures at 30 °C yielded maximal FORC inhibition, providing the basis for transcriptomic and metabolomic analyses to elucidate the biosynthetic mechanisms underlying this inhibition.

**Figure 1.**
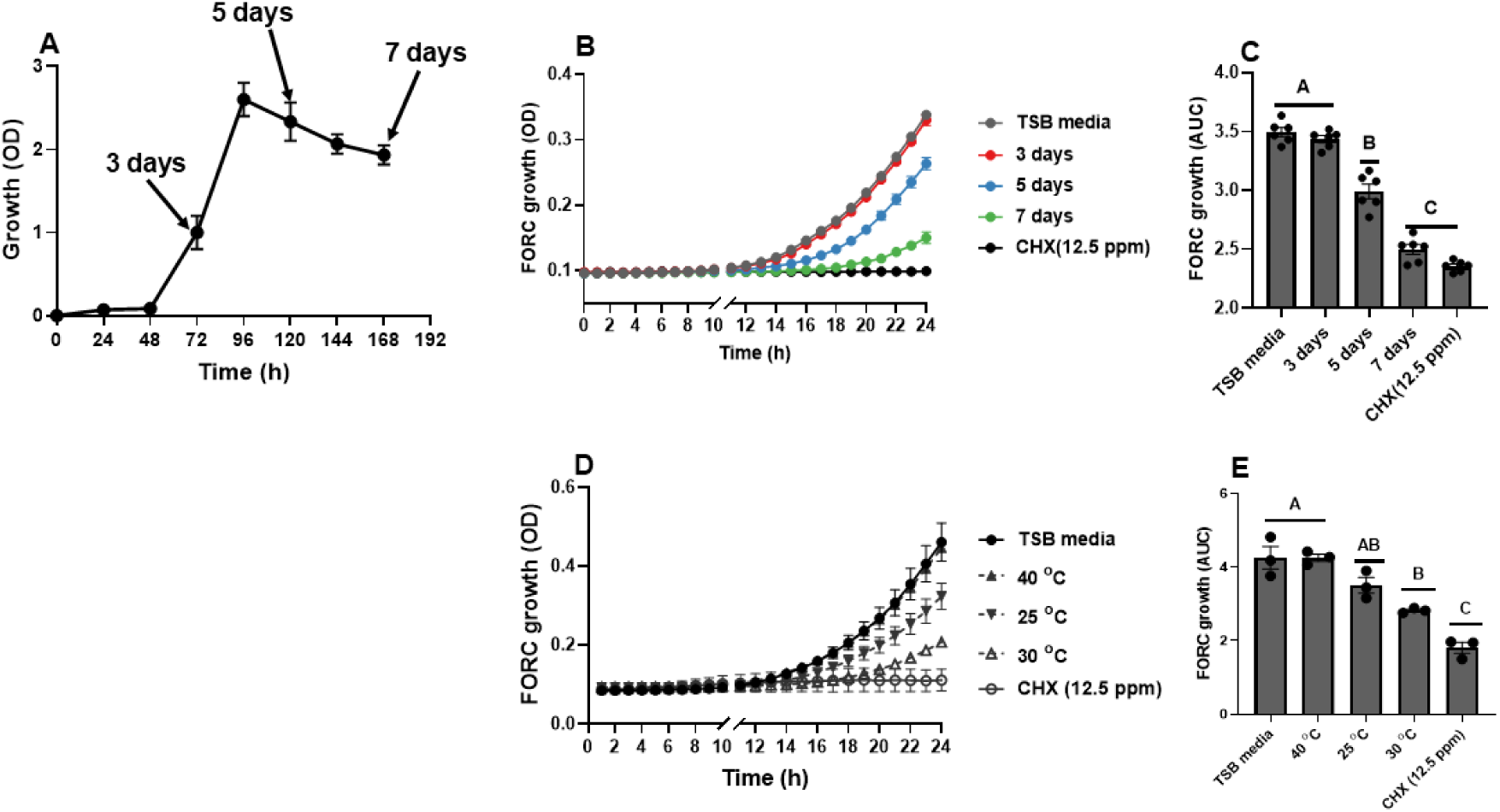
Maximal inhibition of FORC by *S. xinjiangensis* XJ-54 CFS occurs after 7 days at 30 °C of *S. xinjiangensis* XJ-54. Growth curve of XJ54 in TSB medium **(A)**; growth kinetics and area under the growth curve (AUC) of FORC treated with CFS collected at different growth stages **(B-C)** and incubation temperatures **(D-E)** of XJ-54. Growth of FORC was quantified by changes in optical density at 600 nm (OD₆₀₀) monitored over 24 h and normalized to initial readings before estimating the area under the growth curve (AUC). Different letters indicate statistically significant differences among treatments (Wilcoxon test, *P* < 0.05). Data are presented as mean ± standard error of the mean (SEM); data points represent averages of six replicates for five and three independent experiments in panels (**B-C**) and (**D-E**), respectively. TSB medium and cycloheximide (CHX, 12.5 ppm) were used as negative and positive controls, respectively.

### Growth phase facilitates distinct *S. xinjiangensis* XJ-54 transcriptional patterns

To determine which genes and pathways are differentially regulated during the transition to stationary phase, when antifungal activity was highest, we compared transcriptomic profiles of XJ-54 cells collected on days 3, 5, and 7.

*S. xinjiangensis* XJ-54 exhibited distinct transcriptomic profiles across the three time points, with a pronounced shift observed during the late stationary phase (day 7) (**Figure S1A**). Of the 4,343 predicted genes in the *S. xinjiangensis* XJ-54 genome, 2,410 were significantly differentially expressed (log₂FC ≥ 2, adjusted *P* < 0.05) across the time course. K-means clustering of the RNA-seq data identified four distinct gene expression patterns (**Figure S1B**). Cluster K1 (973 genes) comprised genes showing progressive upregulation from day 3 to day 7, whereas Cluster K2 (891 genes) exhibited continuous downregulation across the time points. Cluster K3 (399 genes) displayed high expression on day 3 followed by a marked decline at later stages, while Cluster K4 (147 genes) showed reduced expression on day 5 with partial recovery on day 7.

Functional enrichment analysis using KOBAS was performed on the subset of differentially expressed genes with functional annotations. Of the 4,343 predicted genes, 2,008 were assigned to KEGG or GO terms, of which 1,117 were differentially expressed (log₂FC ≥ 2, adjusted *P* < 0.05) and subjected to enrichment analysis. Cluster K2, comprising genes highly expressed during the logarithmic phase, was substantially enriched in pathways related to primary metabolism and biomass accumulation (**Figure S2**). Translation-associated functions were among the most significantly enriched, including the KEGG Ribosome pathway (adjusted *P* = 5.3 × 10⁻⁸) and GO terms related to structural ribosomal components, indicating elevated protein synthesis during exponential growth. Energy generation pathways were similarly enriched, particularly oxidative phosphorylation (adjusted *P* = 4.4 × 10⁻⁹) and associated aerobic respiration complexes, reflecting the high ATP demand characteristic of rapid cell division. Cluster K3, characterized by high expression between day 3 and 5 followed by a marked decline, showed enrichment of pathways involved in glycine, serine and threonine metabolism, as well as the GO term positive regulation of DNA-templated transcription, while no pathways were significantly enriched in Cluster K4 (**Figure S2**). In contrast, Cluster K1, comprising genes whose expression increased progressively and peaked during stationary phase, displayed significant overrepresentation of fatty acid biosynthesis (adjusted *P* = 5.97 × 10⁻⁵) and metabolism (adjusted *P* = 1.54 × 10⁻⁴), amino acid-related pathways, including alanine, aspartate and glutamate metabolism (adjusted *P* = 1.5 × 10⁻⁴) and nitrogen metabolism (adjusted *P* = 4.6 × 10⁻³) (**Figure S2**).

### BGC expression in *S. xinjiangensis* XJ54 is growth phase-dependent

To identify which BGCs are transcriptionally activated under conditions that yield FORC inhibition (stationary phase, 30°C), we compared expression of all twelve BGCs across the three timepoints. Transcriptional activity of BGCs was quantified as the mean number of transcripts per million (TPM) of differentially expressed core biosynthetic genes within each cluster. The twelve identified BGCs exhibited substantial heterogeneity in both transcript abundance and expression timing (**Figure. 2**). The most highly transcribed clusters were Region 10 (ectoine) and Region 1 (NRP-metallophore/NRPS), which reached peak transcript levels exceeding 60,000 TPM at 7 days and 4,000 TPM at 3 days, respectively.

**Figure 2.**
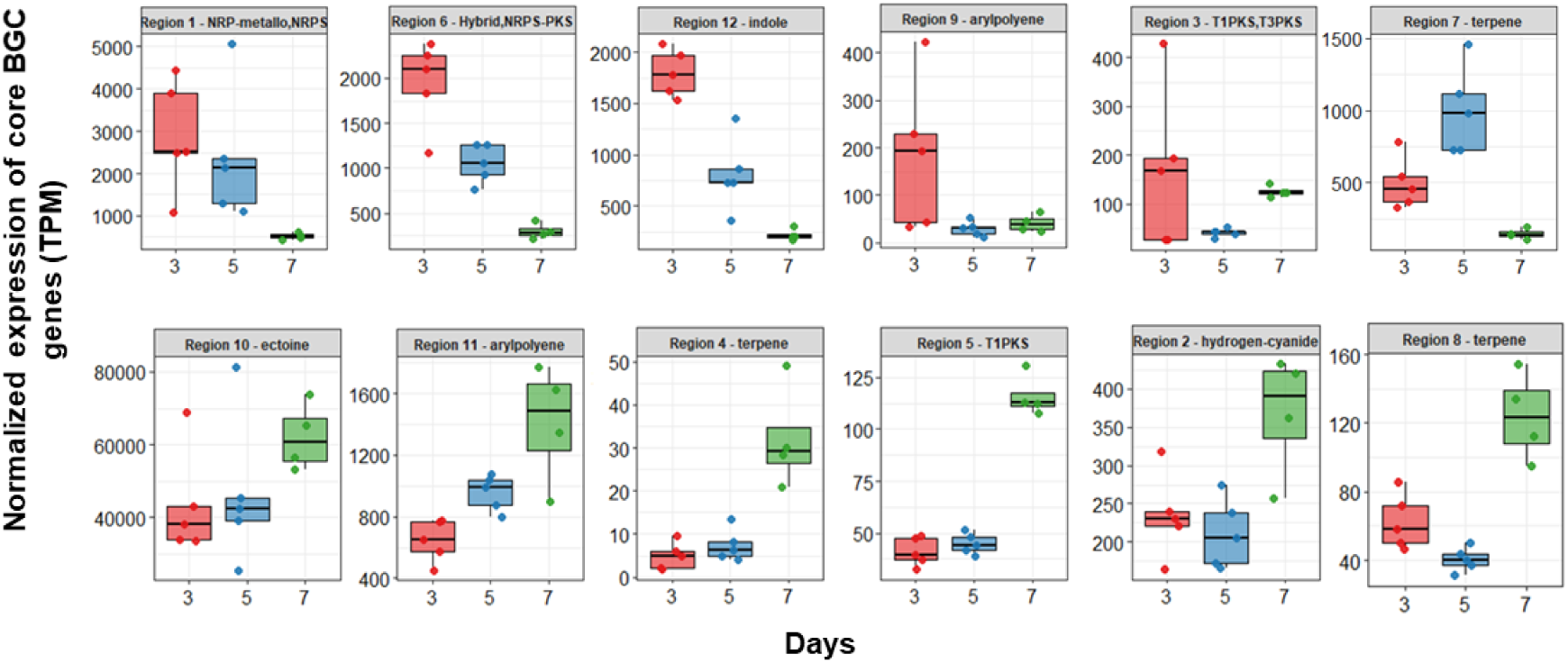
BGC expression in *S. xinjiangensis* XJ-54 is temporally regulated. Normalized expression (transcripts per million, TPM) of core biosynthetic genes for twelve BGCs at 3, 5, and 7 days of growth. Each boxplot represents the mean expression of differentially expressed (log₂FC > 2, adjusted *P* < 0.05) core biosynthetic genes within each BGC in 5 replicates.

Several BGCs were highly expressed during exponential growth phase compared to stationary phase, including Region 1 (NRPS-metallophore), Region 6 (hybrid NRPS-PKS), Region 12 (indole), Region 9 (arylpolyene), and Region 3 (T3-T1PKS). In contrast, Regions 10, 11, 4, 8, 5, and 2, putatively encoding ectoine, arylpolyene, terpenes, T1PKS, and hydrogen cyanide biosynthesis, respectively, were upregulated during stationary phase, and most significantly at day 7. These data revealed pronounced differences in both the intensity and timing of BGC transcription, reflecting dynamic regulation of specialized metabolite biosynthesis throughout the growth of *S. xinjiangensis* XJ-54. The stationary-phase upregulation of Regions 10, 11, 4, 8, 5, and 2 suggests these clusters may encode antifungal metabolites, as this expression pattern temporally correlates with the observed increase in FORC inhibition.

### Transcriptional coupling between regulatory and biosynthetic genes varies temporally across BGCs

Understanding the regulatory mechanisms controlling BGC expression may enable targeted enhancement of secondary metabolite production, particularly the antifungal metabolites that accumulate during stationary phase. To assess the regulation of secondary metabolite biosynthesis, we examined the temporal dynamics of differentially expressed cluster-situated regulatory genes (CSRs) linked to expression of BGC-core biosynthetic genes from the twelve *S. xinjiangensis* XJ-54 BGCs across the three timepoints.

The expression of CSRs and core genes exhibited BGC-specific transcriptional relationships, with substantial variance even among BGCs belonging to the same biosynthetic class, indicating diverse regulatory strategies (**Figure S3**). We identified three distinct regulatory patterns. In Region 10 (ectoine), CSRs were co-expressed with core biosynthetic genes, exhibiting similar temporal transcription profiles. Regions 1 (NRP-metallophore-NRPS) and 8 (terpene) showed a similar transcription pattern although their CSRs were not significantly differentially expressed. Unlike Regions 10, 1, and 8, Region 12 (indole) displayed early co-expression between CSRs and core genes, followed by divergence at later time points. In Regions 4 (terpene) and 11 (arylpolyene), CSRs and core genes displayed negatively correlated temporal expression patterns, with CSRs transcript levels decreasing as core biosynthetic gene expression peaked on day 7, indicative of stationary-phase induced de-repression. Region 2 exhibited a similar inverse correlation, although its CSR was not differentially expressed across time points.

More complex regulatory dynamics were observed in Regions 3 (T1-T3 PKS), 5 (T1PKS), 6 (hybrid NRPS-PKS), 7 (terpene), and 9 (arylpolyene), where CSR and biosynthetic gene expression patterns were heterogeneous. Within these individual BGCs, some CSRs showed synchronized expression with core genes while others exhibited asynchronous dynamics, highlighting the multilayered regulatory architecture governing these clusters.

Among all BGCs, Region 6, a large (∼168 kb) hybrid NRPS-PKS cluster, exhibited the most complex regulatory architecture. It encodes ten CSRs, five of which were differentially expressed and showed mixed co-expression and asynchronous temporal dynamics relative to core biosynthetic genes (**Figure 3**). Two putative activators (a *CheY*-like response regulator and *AIIS*) were highly expressed at day 3, consistent with early cluster activation, whereas two *TetR* family repressors and a *CynR* family regulator were weakly expressed at days 3 and 5 but strongly upregulated at day 7. This late-phase induction of predicted repressors coincided with sustained downregulation of biosynthetic genes, suggesting a shift from early activation to late repression of this BGC.

**Figure 3.**
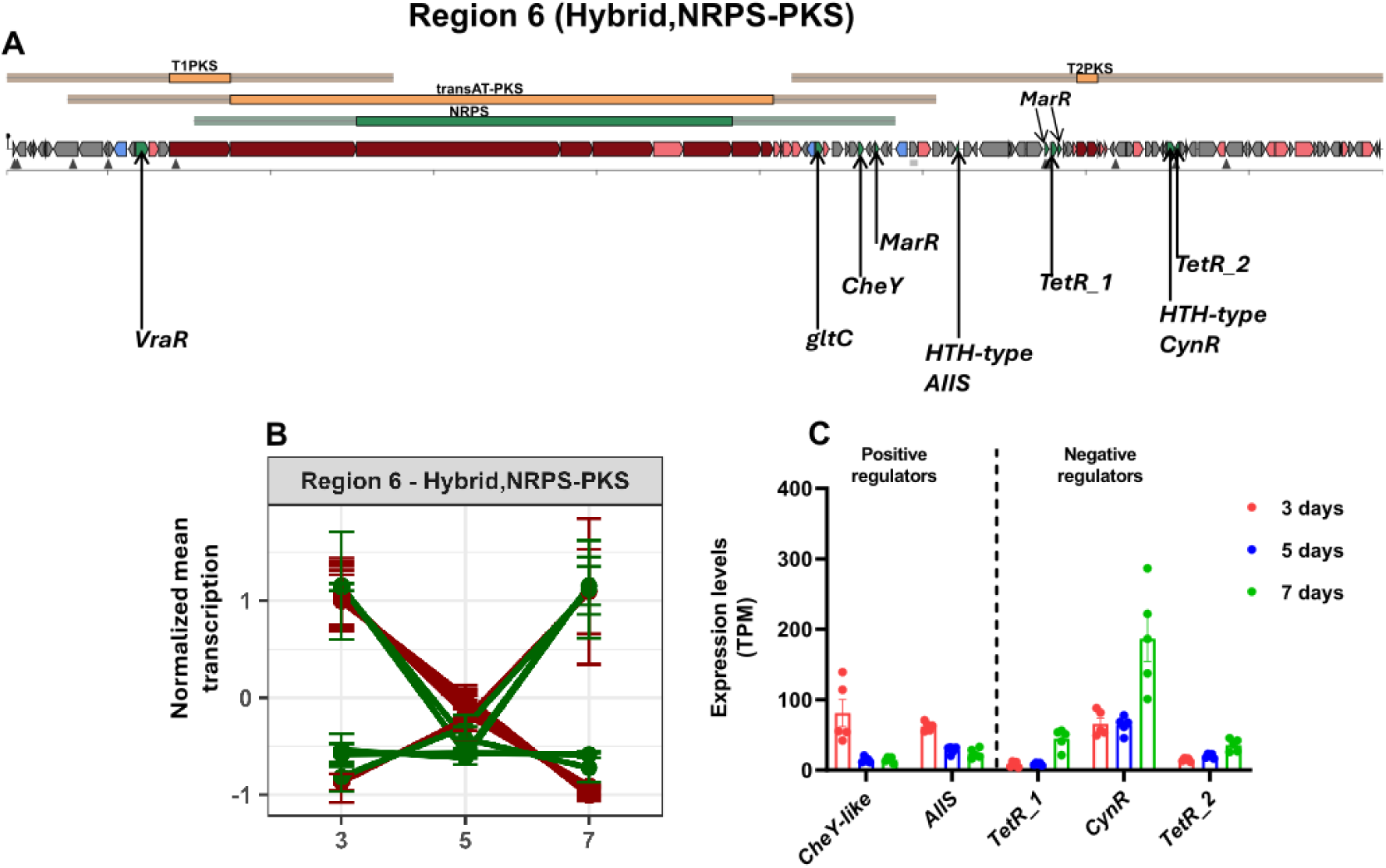
Region 6 (hybrid NRPS-PKS) displayed more complex temporal expression relationships between CSRs and core biosynthetic genes than any other BGC region in *S. xinjiangensis* XJ54. Genomic organization of Region 6 showing predicted biosynthetic modules and cluster-situated regulators (CSRs) **(A)**; Normalized mean transcription of core biosynthetic (red) and regulatory (green) genes at 3, 5, and 7 days **(B)**; Expression profiles (TPM) of differentially expressed CSRs showing early induction of activators and late upregulation of repressors **(C)**.

Overall, CSR-core gene expression relationships were strongly BGC-specific and time-dependent, ranging from coordinated temporal expression to inverse or heterogeneous dynamics, with the greatest complexity observed in Region 6 (hybrid NRPS-PKS).

### Metabolomic profiling of *S. xinjiangensis* reveals growth phase-dependent shifts in metabolite composition and chemical class abundance

To identify the metabolites responsible for the observed antifungal activity, we profiled the extractable metabolome at days 3, 5, and 7, correlating metabolite accumulation patterns with the temporal increase in FORC inhibition. Following removal of media-derived and single-sample features, 255 unique sample-associated metabolite features were retained for analysis. Principal component analysis (PCA) revealed distinct metabolic profiles across time points, with significant separation between day 3 (exponential phase) and day 7 (late stationary phase) (PERMANOVA, adjusted *P* = 0.021; **Figure 4**).

**Figure 4.**
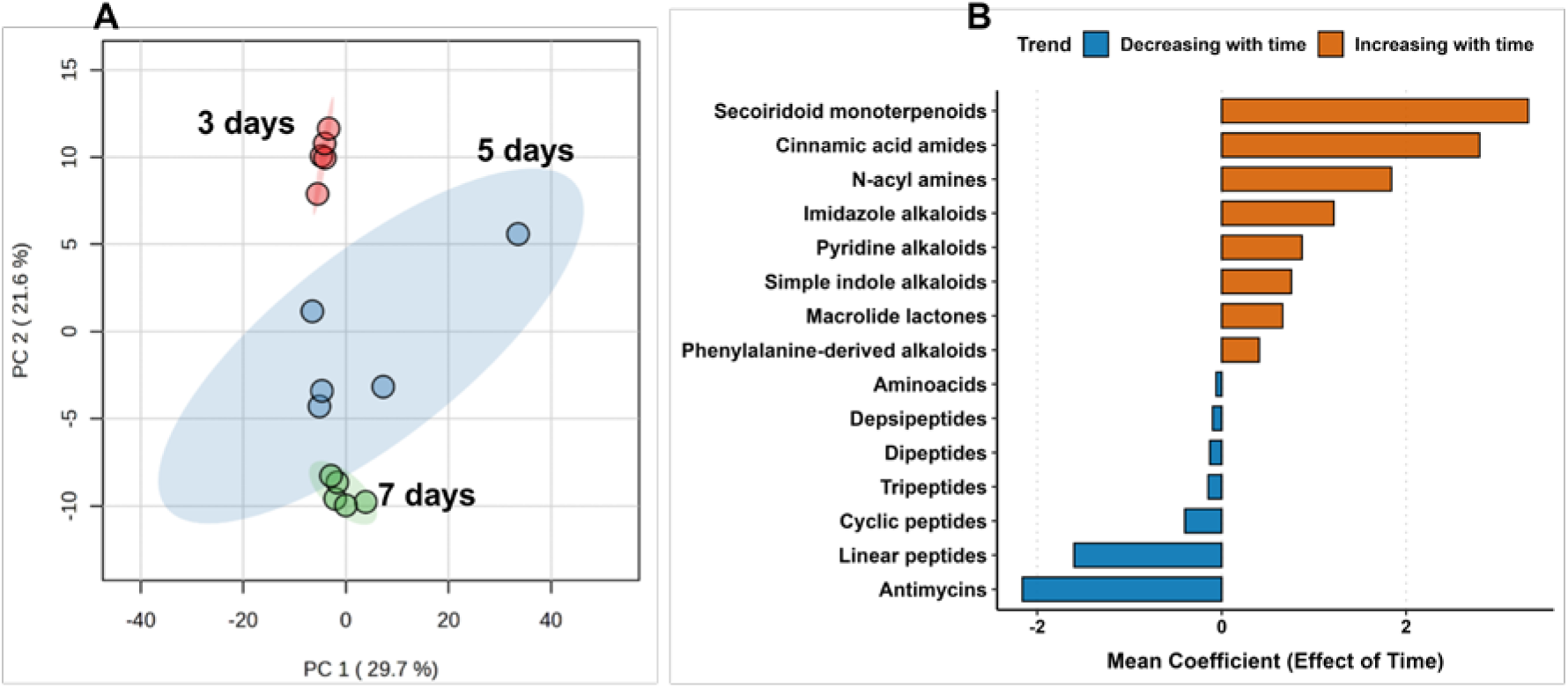
Temporal variation in metabolite composition and class abundance during growth of *S. xinjiangensis*. **(A)** Principal component analysis (PCA) of metabolomic profiles showing distinct clustering of samples from 3-, 5-, and 7-day cultures. Metabolite composition at days 5 and 7 differed significantly from day 3 (PERMANOVA, adjusted *P* = 0.01). **(B)** Mean MaAsLin2 coefficients for significantly changing metabolite classes (adjusted *P* < 0.05). Bars indicate metabolite classes that increased (orange) or decreased (blue) in relative abundance over time.

A feature-based molecular network of all detected metabolites was constructed using the GNPS platform and annotated with NPC pathway predictions from SIRIUS, allowing classification of features into broad metabolite classes. The largest and most interconnected clusters were enriched in amino acids and peptides (68.6%), while smaller subnetworks contained alkaloids (5.5%), polyketides (2.4%), fatty acids (1.2%), carbohydrates (0.8%) and terpenoids (0.4%) and were detected at low frequencies. A substantial fraction of metabolites was unclassified (21.2%), representing chemical entities that are either poorly covered in reference databases or potentially novel. Linear modeling with MaAsLin2 was used to identify metabolites whose abundances changed significantly over time, and the analysis revealed significant time-associated changes in metabolite class composition across the 3-, 5-, and 7-day cultures (adjusted *P* < 0.05) (**Figure 4**). Of the 117 metabolites that changed significantly, 62 increased in abundance over time, while 55 decreased. Grouping these metabolites by chemical class using SIRIUS revealed growth phase-specific production patterns. Secoiridoid monoterpenoids, cinnamic acid amides, N-acyl amines, and several alkaloid classes (including imidazole, pyridine, and indole alkaloids) increased in abundance from the exponential to the stationary phase, whereas antimycins, linear peptides, and cyclic peptides decreased.

Together, these results reveal distinct temporal patterns in metabolite class abundance, with specific chemical groups enriched or depleted across different growth stages. The accumulation of alkaloids, N-acyl amines, and monoterpenoids during stationary phase (on day 7), coinciding with peak antifungal activity, suggests these metabolite classes contribute to FORC inhibition.

### Bioassay-guided evaluation of CFS fractions uncovers potent antifungal activity and hyphal damage associated with fraction 19

To prioritize purification efforts and pinpoint metabolites contributing to antifungal activity, CFS collected at day 7 were fractionated using flash chromatography, yielding 19 fractions that were tested for antifungal activity. Each fraction was reconstituted in DMSO, diluted in sterile distilled water, and tested against FORC spores (10⁵ spores mL⁻¹) in 96-well microplate assays (final DMSO concentration, 0.05%). PDB medium, DMSO (1%), and cycloheximide (12.5 ppm) served as negative, solvent, and positive controls, respectively. Only fraction 19 consistently inhibited fungal growth across three independent experiments (**Figure. 5A and Figure S4**). Microscopic examination of FORC spores treated with this fraction for 18 h revealed hyphal damage, as evidenced by propidium iodide and calcofluor white staining (**Figure. 5C, bottom**). In silico structural annotation of fraction 19 based on LC-MS/MS data revealed a main peak of m/z 257.04776 with an isotopic pattern corresponding to the molecular formula C₁₄H₉ClN₂O, suggesting a halogenated alkaloid (**Figures S5-S8**). The identification of a halogenated alkaloid in the bioactive fraction is consistent with the metabolomic observation of alkaloid accumulation during stationary phase, when antifungal activity was maximal.

**Figure 5.**
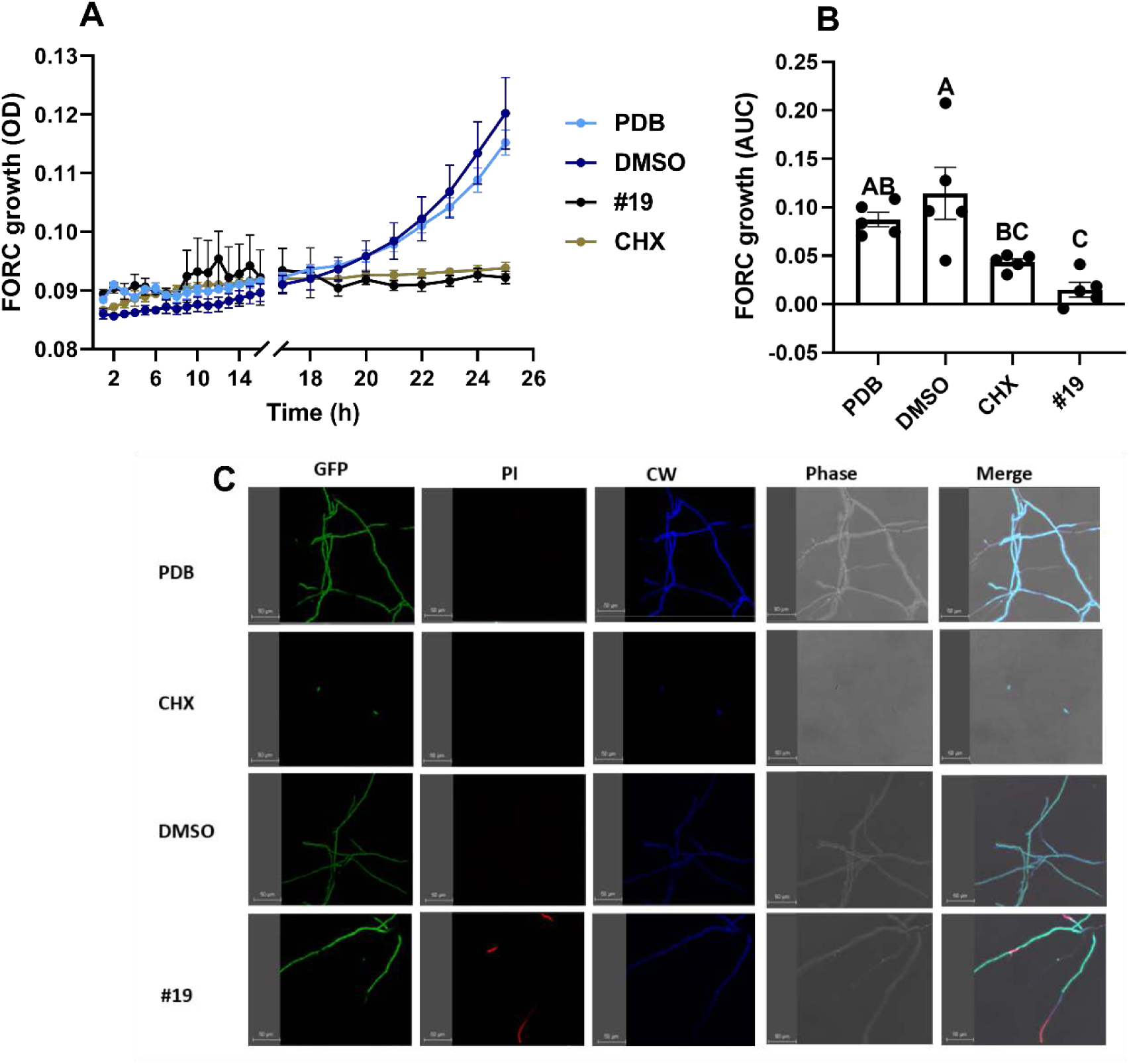
Growth kinetics **(A)** and area under the growth curve **(B)** of FORC treated with fraction 19 in in-vitro 96-well plate experiments. Fraction 19 exhibited clear FORC growth inhibition compared to negative (PDB medium), solvent (DMSO, 1 %), and positive (CHX; cycloheximide, 12.5 ppm) controls. Different letters indicate statistically significant differences among treatments (ANOVA, *P* < 0.05). Data represent means ± SEM (n=5). Microscopic visualization of GFP-FORC spores treated with fraction 19 for 18 h and stained with propidium iodide (PI) and calcofluor white (CW) **(C)**. Red-stained hyphae and disrupted calcofluor white-stained walls (bottom panel), indicative of hyphal cell wall damage. Images were obtained by confocal microscopy.

## Discussion

Soil-derived actinomycetes harbor extensive biosynthetic potential encoding antifungal secondary metabolites, yet natural product discovery has remained disproportionately focused on *Streptomyces*, leaving the metabolic capabilities of rare actinomycete genera such as *Saccharomonospora* uncharacterized (10, 11). While genome mining has revealed extensive BGC repertoires in microbial genomes, relatively few studies have directly examined BGC expression in native hosts or under defined culture conditions (14, 50), despite frequent assertions that most BGCs are transcriptionally silent under laboratory conditions. Understanding the conditions that trigger BGC expression and secondary metabolite accumulation is critical for optimizing bioactive compound production. This study aimed to elucidate mechanisms underlying the temporal antifungal activity of a rare actinomycete, *S. xinjiangensis* XJ54 through application of an integrated omics approach.

In vitro bioactivity assays revealed that *S. xinjiangensis* XJ-54 produced antifungal metabolites that inhibited the growth of the phytopathogen FORC exclusively during stationary phase. These findings are consistent with studies in *Streptomyces* showing that the transition from late exponential to stationary growth is critical for activating antifungal secondary metabolites (51). The observed temporal regulation reflects the well-established physiology of secondary metabolism in Actinobacteria. When nutrients become limiting during the transition from exponential to stationary phase, many actinomycete bacteria, including *Saccharomonospora,* initiate a complex developmental program directed toward sporulation (52). This transition has been linked to the production of bioactive secondary metabolites with antibiotic, antifungal, antiviral, antitumor, or insecticidal activities (52, 53). Additionally, changes in the culture physical conditions accompanying this transition may further modulate metabolite biosynthesis. During microbial fermentation, accumulation of metabolic by-products commonly drives pH fluctuations that can trigger or enhance antimicrobial production (54, 55). We observed a notable rise in medium pH in XJ-54 batch cultures (from 7.6 during exponential growth to 9.0 in stationary phase). Such alkaline pH shifts have been reported to stimulate secondary metabolism in *Streptomyces*, for instance, increasing production of validamycin A (56) and actinorhodin (57), suggesting that the observed pH increase may contribute to induction of antifungal metabolite biosynthesis.

While it is often reported that most BGCs are transcriptionally silent under standard laboratory conditions (13), we observed that 68% of biosynthetic genes across the 12 BGCs in *S. xinjiangensis* XJ-54 were significantly expressed, indicating that transcriptional inactivity is not a universal feature of BGCs. This is consistent with transcriptomic analyses in *Salinispora* spp. (15), myxobacteria (58), and *Streptomyces* (51), which similarly exhibited widespread BGC expression under routine cultivation. Our results further reveal that 50% of the BGCs were highly expressed during exponential growth and 50% during stationary phase, despite studies citing that BGC expression is often linked to nutrient limitation during the onset of stationary phase (55). Comparable growth-phase dependent patterns have been reported in *Salinispora* (15), myxobacteria (58), and *Streptomyces* (51). Together, these observations suggest that early-expressed BGCs may contribute directly to growth-related physiological processes, whereas those induced later may function in stress adaptation.

The onset and magnitude of antibiotic biosynthesis are controlled by multilayered regulatory networks acting at hierarchical levels. At the pathway level, cluster-situated regulators (CSRs) directly modulate transcription of adjacent biosynthetic genes, allowing individual BGCs to respond to physiological states and environmental cues with precise temporal control (59, 60). Temporal comparison of CSR and core biosynthetic gene expression revealed that BGC regulation is highly cluster-specific and time-dependent, with BGCs exhibiting diverse CSR-core gene relationships, ranging from coordinated expression, inverse temporal dynamics, and heterogeneous patterns, especially in BGCs encoding more than one CSR. The large hybrid NRPS-PKS Region 6 exhibited the highest regulatory complexity, with early expression of putative activators followed by later induction of predicted repressors coinciding with reduced transcription of core biosynthetic genes.

The tight regulatory coordination likely reflects an evolutionary strategy to minimize the energetic cost of specialized metabolite production, ensuring biosynthesis occurs only when timely and appropriate (60). In *Bacillus subtilis*, NRPS/PKS antibiotic production was shown to incur a fitness cost, manifested as reduced growth, and was conditionally induced in response to specific cues, reflecting fitness trade-offs that restrict costly biosynthetic expression to ecologically relevant contexts (61). Because CSRs act as proximal control points for BGC expression, identifying the growth phases and culture conditions that induce their transcription provides a rational basis for BGC prioritization, experimental validation, and targeted induction of specialized metabolite production.

Time-resolved metabolomic profiling revealed pronounced growth-phase-associated shifts in the detectable metabolite pool of *S. xinjiangensis* XJ-54. As cultures entered stationary phase, a subset of annotated features assigned to secoiridoid-like monoterpenoids, N-acyl amines, and alkaloids (imidazoles, pyridine and indoles) increased in relative abundance. These classes of alkaloids have been previously isolated from diverse actinomycete groups and exhibit a broad range of bioactivities such as antibacterial, antifungal, and cytotoxic activity (62). Bioactivity-guided fractionation identified a fraction from stationary phase crude extracts with antifungal activity and LC-MS/MS analysis of this fraction revealed a dominant ion displaying an isotopic pattern consistent with chlorination, and in-silico annotation suggested an alkaloidal scaffold.

Chlorinated alkaloids and other halogenated nitrogen-containing metabolites are well documented among microbial natural products produced by actinomycetes, and are frequently reported as antimicrobial agents (63). Halogenated indole alkaloids were obtained by co-culturing the two actinomycetes *Saccharomonospora* sp. UR22 and *Dietzia* sp. UR66 (64). Alkaloids serve multiple roles in producer organisms, including competitor inhibition and cell-to-cell communication, and have been shown to act as signaling molecules in bacteria (65, 66). However, from an applied perspective, alkaloids have been increasingly reported to exhibit antifungal activity against phytopathogenic fungi, prompting interest in their potential relevance to crop protection (67).

In an attempt to link the BGCs identified via transcriptomics to the active compounds found in the metabolomic analysis, we identified two transcriptional features that likely drive the stationary-phase accumulation of alkaloidal metabolites in *S. xinjiangensis* XJ-54. First, we identified genes involved in alanine, aspartate, glutamate, and nitrogen metabolism that were significantly upregulated during stationary phase, consistent with nitrogen recycling necessary for providing precursor pools for alkaloid biosynthesis (68). Second, *S. xinjiangensis* XJ-54 harbors an indole-encoding BGC that could directly participate in indole alkaloid biosynthesis (68). Although this BGC showed peak transcription during logarithmic growth rather than stationary phase, translation of secondary metabolic genes has been previously shown to correlate poorly with transcript levels (69), suggesting that BGC products may accumulate despite reduced transcript abundance during stationary phase.

Importantly, the majority of detected metabolites could not be confidently assigned to known chemical classes, reflecting both limited spectral library coverage and the chemical novelty typical of rare actinomycetes. Moreover, the extracted metabolome may not fully represent *S. xinjiangensis* XJ-54’s chemical output, as solvent choice, extraction conditions, and ionization bias can preferentially enrich certain metabolite classes while excluding others. Consequently, the annotated metabolites likely represent only a small, biased fraction of the true metabolome.

Overall, these results indicate that antifungal activity in *S. xinjiangensis* XJ-54 is temporally associated with stationary-phase metabolite production, and that an antifungal fraction containing a putative chlorinated alkaloid contributes to this activity. Further purification and NMR structural validation will be required to pinpoint the active compound and resolve its structure. While many BGCs were differentially expressed across the growth stages of *S. xinjiangensis* XJ-54, we could not directly link them to their cognate metabolites, as nearly all of these clusters encode currently uncharacterized products. Nevertheless, the temporal transcription data provides a starting point for BGC prioritization and functional characterization in future experimental studies. More broadly, the identification of antifungal metabolites from rare actinomycetes like *S. xinjiangensis* XJ-54 addresses a critical need in both agriculture and medicine. The emergence of fungicide-resistant plant pathogens threatens global food security, while the rise of drug-resistant fungal infections poses increasing clinical challenges. Underexplored actinomycete lineages, with their distinctive biosynthetic potential, offer opportunities to discover novel scaffolds that may serve as leads for next-generation antifungal development.

